# Mutational scan of self-cleavage by HIV-1 protease provides new views of a conformationally dynamic mechanism

**DOI:** 10.1101/2025.10.07.680924

**Authors:** Gily S. Nachum, Jeffrey I. Boucher, Julia M. Flynn, Princess Quansah, Sivan Nachum, Daniel N.A. Bolon

**Affiliations:** Department of Biochemistry and Molecular Biotechnology University of Massachusetts Chan Medical School, 364 Plantation St, Worcester, MA 01605 USA

## Abstract

The mechanism of conformationally dynamic proteins remains understudied because they are difficult to analyze structurally. For HIV-1 protease the mechanism of cleavage by mature protease is well understood in large part because it forms a stable structure that is amenable to x-ray crystallography. However, self-cleavage or autoproteolysis of protease from the viral polyprotein involves transiently populated structures and is poorly understood. We probed autoproteolysis in HIV-1 using a yeast reporter and mutational scanning. We compared our results with mutational scanning of protease on viral fitness, which integrates both autocleavage and cutting by mature enzyme. We identified 220 mutations that were well tolerated for self-cleavage but not fitness. We analyzed three of these mutations (D30E, W42M and P44L) using independent approaches. All three were capable of efficient self-cleavage in a bacterial assay of autoproteolysis, but had strong defects in mature form for cleavage of a peptide substrate. These separation of function mutations from the mutational scan clustered at hot-spot locations that do not impact autoproteolysis likely because they are conformationally dynamic during self-cleavage. We used the mutational scanning results and molecular simulations to provide models of autoproteolysis conformations. These models provide new views of a structurally dynamic mechanism.

## Introduction

Dynamic motions of proteins are critical to function (1–8), but the mechanism of most structurally dynamic proteins are poorly understood because dynamics are more challenging to elucidate than structures of stable states (9). Nuclear magnetic resonance (NMR) approaches have been developed to provide direct readouts of motions over a large range of timescales(9), but cannot be efficiently applied to some proteins due to their size or solubility. For example, the mechanism of autoproteolysis (AP) of HIV-1 protease (PR) involves a transiently populated state in a large polypeptide making it virtually impossible to investigate by high-resolution physical approaches. While decades of intensive research into HIV-1 has provided detailed structural and mechanistic insights into most aspects of viral infectivity, key pieces of the lifecycle remain poorly resolved – particularly those that involve dynamic conformations including autoproteolysis.

The genome of the HIV-1 retrovirus encodes two polyproteins, Gag and Gag-Pol, with a ribosomal frameshift occurring in approximately 5% of the translational events resulting in the Gag-Pol polyprotein (10, 11). These polyproteins must be cleaved into individual proteins by PR in order to generate infectious virions (12–15). PR is initially synthesized as part of this Gag-Pol polyprotein and is an essential enzyme in the viral life cycle, functioning as a 99-residue symmetric dimer with a single active site with catalytic residues contributed by each subunit (16–18). The mature protease dimer is composed of β-strands and flexible flaps that adopt open and closed conformations with the help of hinge domains (19, 20). These flaps open to allow substrate entry into the binding pocket and then close around the substrate to position it near the protease’s two catalytic aspartate residues. Mature PR recognizes and cleaves 12 defined sites in the Gag and Gag-Pol polypeptides with diverse amino acid sequences, demonstrating both flexibility and specificity (21).

Following translation, PR is flanked by p6* at its N-terminus and reverse transcriptase (RT) at its C-terminus. This precursor-PR must first cleave itself out of the polyprotein at its N- and C-terminal ends in a process termed autoproteolysis (AP) during which precursor-PR serves as both substrate and enzyme. Interestingly, AP does not occur immediately after translation of the Gag-Pol but is temporally coordinated with the release of the budding viral particles by an unknown mechanism. Premature PR activation results in processing of viral proteins prior to virion formation and inhibits viral budding (22). Many studies of AP have utilized minimal constructs containing PR along with upstream and downstream peptides of less than 100 amino acids from Gag-Pol (23–28). Mutation of the catalytic aspartate (e.g., D25A) abolishes AP activity, suggesting that the enzyme mechanism of cleavage is largely shared with that of mature PR (29). AP in these contexts is concentration dependent, indicating that AP requires dimerization (23). An intramolecular cleavage of a transient dimeric species of PR at the N-terminal p6*–PR junction has been found to be essential for AP (30). Mutations that prevent cleavage at this site result in an inactive protease precursor, a defect in Gag polyprotein processing, and non-infectious viruses (31). Furthermore, four amino acid extensions at the N-terminus of mature protease have been shown to disrupt the β-sheet that stabilizes dimerization, dramatically reducing enzyme activity (23). In contrast, extensions at the C-terminus of mature protease do not affect dimerization or enzymatic activity. Kinetically, the cut site at the N-terminus of PR is processed faster than the site at the C-terminus during AP. Analyses of full-length Gag-Pol have shown similar AP patterns (30) indicating that the minimal constructs report on biologically relevant mechanisms.

Despite sharing similar mechanisms and identical amino acid sequences, evidence suggests that the structures of mature PR and precursor-PR may be quite different. In comparison to mature PR, the N-terminal tail of PR must insert itself into the active site in the AP complex (32). Precursor-PR is only able to cleave a small subset of the cleavage sites of mature PR (33). Additionally, precursor-PR is significantly less sensitive to PR inhibitors that bind in the active site(30). While PR is only active as a dimer, it is believed that the upstream p6* peptide significantly destabilizes the dimeric precursor-PR. Thus, precursor-PR is only capable of forming transient, dynamic, low-populated dimers (32), the structure of which remains elusive. Recent cryo-EM analysis of the Gag-Pol polyprotein of HIV-1 shows clear density for RT, indicating that it forms a stable structure, but weak density for other regions prevents a clear understanding of the structure of PR within Gag-Pol (34).

Preventing AP completely disrupts the infection cycle because it prevents the accumulation of mature PR and cutting of multiple other essential sites in the viral polyproteins. This amplified impact of AP on viral infections has made it a desired target for inhibitors. While inhibitors with high affinity for mature PR have been developed, the impacts of these inhibitors on AP are dramatically weaker, typically showing a thousand fold or greater decrease in apparent affinity (35). Motivated by its importance for infection, multiple efforts have been made to elucidate the structure of PR in viral polyproteins(34, 36), but have not yielded clear views likely due to highly dynamic structural properties. Further elucidation of the distinct structure of precursor-PR will provide opportunities to design novel inhibitors that specifically target AP.

To gain insight into the sequence-structure relationship that underlies AP, we performed a comprehensive mutational scan to quantify how all possible point mutations impact AP using a minimal construct (Fig. 1). Similar to how structure-activity relationships illuminate what parts of an inhibitor are critical for binding (37), comprehensive mutational scans provide information on what parts of a proteins are critical or dispensable for function (38–41). Because the readouts of mutational scans are functional, they can be readily applied to any protein independent of its dynamics, solubility, or other physical properties (42, 43). Our analysis of PR revealed key features required for AP function that we used to generate structural models of a highly dynamic yet critical feature of a human pathogen that has been a long-sought goal in virology.

**Figure 1.**
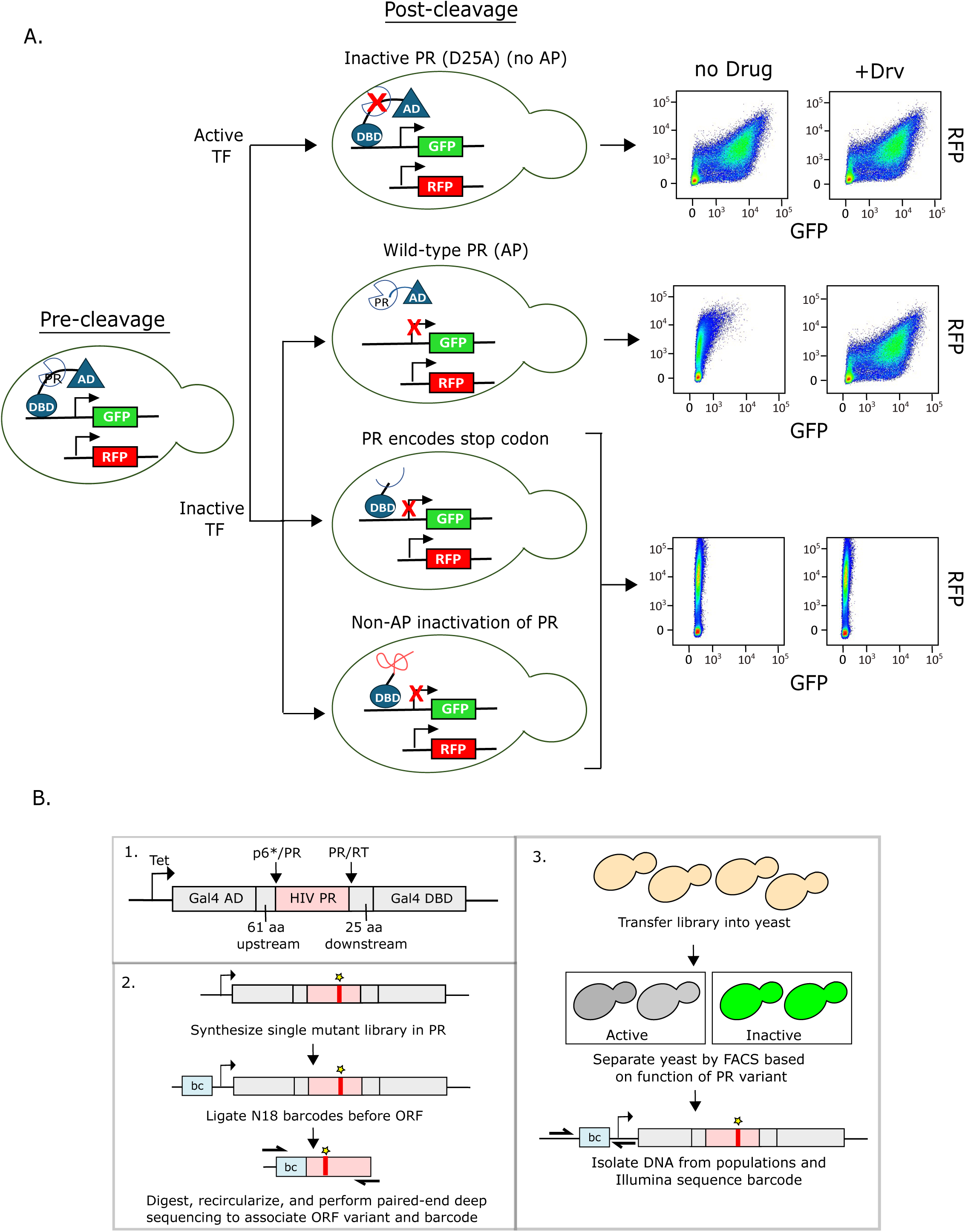
Experimental strategy to measure the function of all individual mutations of HIV protease on autoproteolysis. A. HIV PR was cloned between the DNA binding domain (DBD) and activating domain (AD) of the Gal4 transcription factor. The Gal4 TF drives GFP expression from a galactose promoter. PR variants were sorted based on their ability to cleave themselves out of the TF. Inactive PR (i.e. D25A) is unable to cleave itself out of the TF, resulting in high GFP expression. WT PR successfully cleaves itself out, inactivating the TF, resulting in low GFP expression. Internal stop codons in PR result in truncated and inactive TF. Additionally, certain mutations may inactivate the TF in a non-AP dependent manner, for example by sequestration in aggregates and/or destruction by endogenous yeast enzymes. The PR inhibitor, Darunavir (Drv), specifically inhibits inactivation of the TF in an AP-dependent manner. Cells were separated by FACS based on GFP and RFP expression levels. B. Barcoding strategy to measure frequency of all individual mutations of PR in a single experiment.

## Results and Discussion

To systematically characterize the effects of all protease mutations on AP we developed an efficient mutational scanning approach using a yeast reporter strategy (Fig. 1). To measure AP function, we adapted an approach previously developed by Das Mahaptara and colleagues (44) that used a split transcription factor with PR and flanking regions inserted between the Gal4 DNA binding domain and activation domain. Prior work (44) had used a colorimetric reporter and shown that catalytically dead variants (e.g., D25A) or inhibitors that target PR would cause the transcription factor to remain intact and the reporter active, while wildtype (WT) PR sequences would auto-cleave, leading to a lack of reporter signal. We adapted the approach to a fluorescent readout using a GFP reporter for PR activity and RFP driven by a constitutive promoter to provide a readout of cell size and protein synthesis capacity. WT and catalytically dead D25A PR are clearly differentiated by flow cytometry (Fig. 1A), with WT PR resulting in very low GFP expression and D25A PR having increased GFP expression. High concentrations of the drug Darunavir (Drv) that inhibits PR results in increased GFP expression to similar levels as the inactive D25A mutant, indicating AP can be inhibited (Fig. 1A). The insertion of a premature stop codon in PR leads to a lower level of GFP reporter than WT that is unaffected by Drv. Additionally, we hypothesize that mutations that fully unfold PR may lead to inactive reporter activity that does not result from AP (e.g., sequestration in aggregates and/or destruction by endogenous yeast enzymes). We would expect that variants that lead to an inactive reporter in a non-AP dependent manner would have similar flow cytometry profiles to stop variants (lower level of GFP reporter compared to WT and/or refractory to Drv inhibition) (Fig. 1A). Taken together, these analyses indicated that flow accelerated cell sorting (FACS) would accurately separate PR mutants based on their AP function.

To facilitate the accurate and efficient quantification of all possible point mutations in PR for AP function, we used a barcoding strategy (Fig. 1B). We constructed a library engineered with all possible point mutations in PR into the AP reporter construct and added a random (N_18_) barcode in an inert region of the plasmid. We used a subassembly strategy where paired-end sequencing links each barcode to the PR variant encoded on the same plasmid molecule. This facilitates short read sequencing of barcodes in selection experiments to provide accurate estimates of each PR variant. This approach has three big advantages compared to direct readouts of mutants in mutational scanning experiments: (1) it can track mutants over large stretches of sequence, (2) selection readouts can be analyzed with cost-efficient short-reads, and (3) it reduces the impacts of misreads on readouts of function. The large diversity of possible barcodes (7×10^10^ possible N_18_ combinations) leads to multiple sequence differences between the subset associated with mutant variants. This is important because misreads of a base in a barcode almost never lead to an improper read of another mutant variant. In contrast, misreads from direct sequence readouts frequently lead to improper reads of another mutant variant.

Equipped with these reagents, we determined the impact of each amino acid mutation in PR on its AP function (Fig. 2). The barcoded split Transcription factor (TF) libraries under the control of a Tetracycline (tet)-off promoter were transformed into yeast harboring the GFP reporter in two separate biological replicates. Cells were grown for 48 hours in the presence of tetracycline to inhibit PR expression in order to amplify the libraries without selection. Subsequently, tetracycline was removed to induce PR expression and cells were grown in the absence or presence of the PR inhibitor, Darunavir, for 12 hours. Cells were sorted by flow cytometry into cut (low GFP) and uncut (high GFP) populations (Fig 2A). The sorted populations of yeast were outgrown for an additional 18 hours, plasmids encoding the split TF libraries were recovered, and the barcoded region was amplified by PCR and sequenced using single end Illumina sequencing.

**Figure 2.**
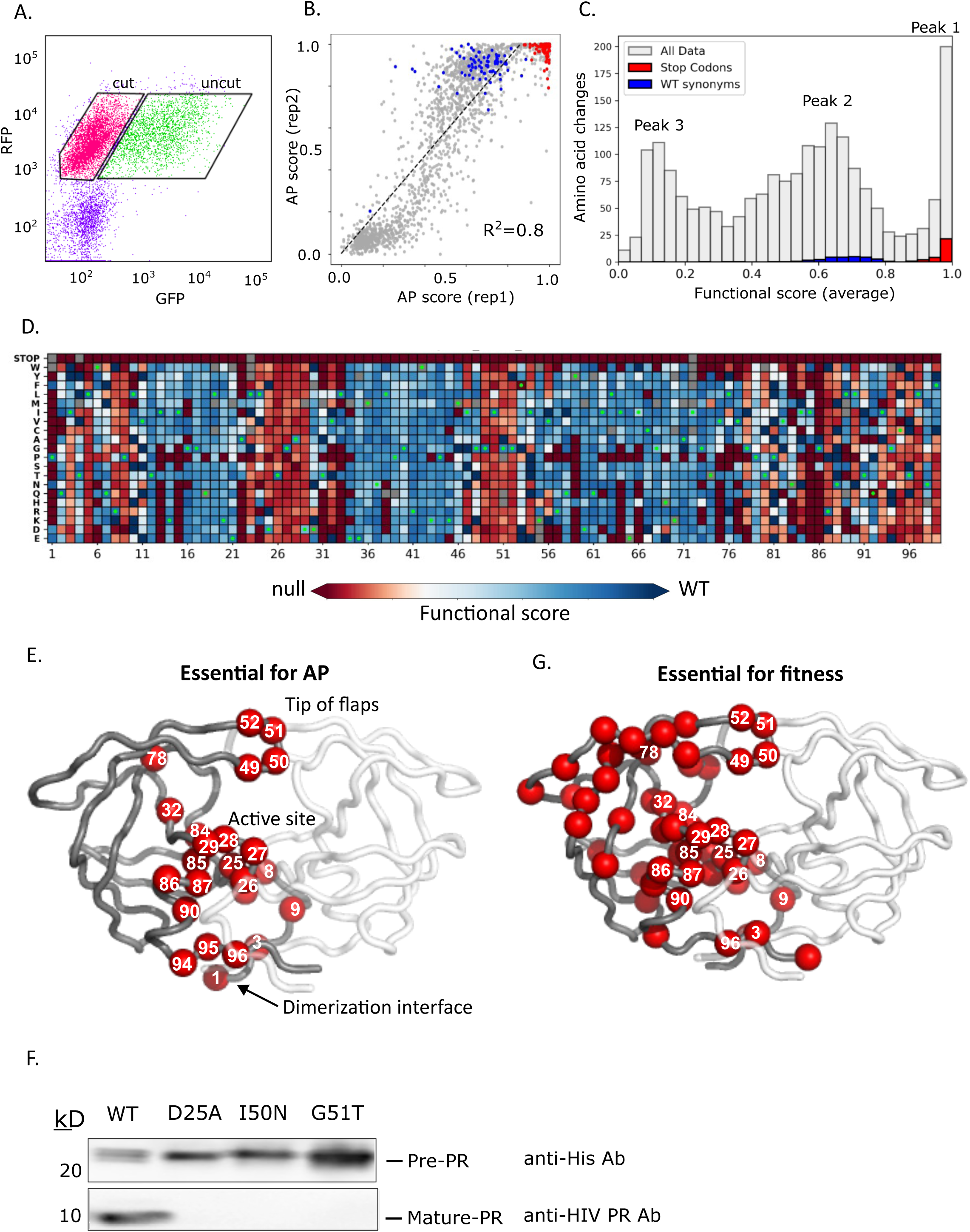
Systematic analyses of the effect of PR mutations on AP function. A) Cells harboring the split-TF PR library were sorted by flow cytometry into cut (low GFP) and uncut (high GFP) populations. B) Correlation between replicates of functional scores of all PR variants between two replicates. Stop codons are shown in red and wild-type synonyms in blue. C) Distribution of all functional scores for all variants (gray), stop codons (red), and wild-type synonyms (blue). Functional scores were calculated as the average between two replicates. D) Heatmap representation of all functional scores of PR. Green dots represent the WT amino acid at each position and grey squares are variants missing from the dataset. E) Positions of PR that are intolerant of mutations for autoproteolysis (15 or more substitutions having null-like function) are represented by red spheres on one subunit. F) Mutations at positions at the tip of the PR flaps that abolish AP in the library screen were measured independently for ability to self-cleave when expressed in bacteria. The PR precursor was monitored with an anti-his antibody against its C-terminal his tag and mature protease with an antibody against the free N-terminus of mature PR. I50N and G51T PR were as defective as the active site mutant D25A in cutting themselves out of the pre-protease construct. G) Positions of PR that were found to be critical for mature protease function as measured by ability of variant to support viral fitness in an infection assay(45) as shown in red. Critical positions are defined as positions with 15 or more mutations causing null-like viral fitness.

We calculated a functional score for each point mutation variant based on its sequencing observations in the cut window relative to both the cut and uncut windows (Table S1). Using this approach, a functional score of 1 indicates a variant that is only observed in the cut window, while a functional score of 0 indicates a variant that is only observed in the uncut window. Functional scores from replicates are strongly correlated (Fig 2B) indicating strong signal to noise in our measurements. Despite the strong correlation, there is a noticeable non-linear relationship between replicates that we suspect is related to differing stringency of selection from slightly different experimental properties between replicates that were performed on different days (e.g., slight differences in expression levels of PR between replicates) (Fig. 2B). For this reason, we calculated the average functional score between the two replicates for each PR variant. The average functional score for members of the library containing silent mutations (WT synonyms) was 0.78, indicating that the WT PR did not cleave the TF to completion. The average functional score for a member of the library containing a stop codon was 0.97, indicative of a fully truncated non-functional TF. Additionally, we calculated a functional score for each variant in the presence of 30 uM Darunavir (Table S1). Drv inhibited WT PR, resulting in an average functional score for WT synonyms of 0.45. For each variant we assigned a score based on how its activity responds to Drv, calculated as the functional score in the absence of Drv divided by the functional score in the presence of Drv (Table S1, Fig. S1A).

The distribution of functional effects of all the split TF library variants in the absence of Drv has three main peaks (Fig 2C). Our analysis indicates that the peak with the highest functional scores within the range of stops and above the range of WT synonyms (labeled Peak 1) are predominantly due to non-AP mechanisms such as aggregation or unfolding making them prone to destruction by yeast proteases. A number of observations support this hypothesis. First, the vast majority of mutants with these high functional scores are not able to support viral fitness (45) (88%, Fig. S1B) indicating that most of these protease variants are inactive. Second, the majority of these variants with high functional scores were non-responsive to Drv (Fig. S1C). The mutants in this group that remain responsive to Drv we hypothesize is due to structural stabilization of PR by the small molecule inhibitor causing them to be refractive to unfolding. Finally, we tested two of these individual mutants (Y38D and Y59S) with high functional scores (FS = 0.95 and 0.96 respectively) for their ability to auto-cleave. To do this, we expressed a PR precursor sequence containing 59 aa upstream and 23 aa downstream of the mutant PR. Self-cleavage activity of the PR precursor was monitored by SDS-PAGE as described in Louis et. al (23). We found that these protease mutants were completely defective for self-cleavage (Fig. S1D). Thus, we conclude that this population of variants with abnormally high functional scores are due to non-AP mechanisms and we classified these as null for AP and assign them a functional score of 0.

The second peak in the distribution of fitness effects in Fig. 2C (Peak 2; FS ∼0.65) overlaps with WT variants indicating that these are highly functional for AP. Concordant with this, the vast majority (>95%) of these variants are responsive to Drv (Fig. S1C). The peak with the lowest FS (Peak 3; FS ∼ 0.1) overlaps with known catalytically dead variants (e.g. mutations to the catalytic D25), indicating that these variants have low to no AP function. From a heatmap representation of the AP results (Fig. 2D), it is clear that there are many positions that are highly sensitive to mutation, indicating that the WT amino acid is critical to structure and/or function. These critical positions, defined as positions with 15 or more substitutions with null-like function (Fig 2E), include the catalytic triad, D25, T26, G27, as well as positions immediately adjacent to the active site. Residues at the dimerization interface (aa 1,3,5 and 94-96) also cannot be altered without complete loss of function, supporting a large body of work previously showing that dimerization is indispensable to AP function (23, 25, 32). Additionally, amino acids 49-52 that are at the tip of a region referred to as “flaps” that can open and close over substrate in mature protease (19, 20) exhibit extremely low mutational tolerance for AP function. To confirm that the WT amino acids at the tips of the flaps are critical for AP function, we tested two individual mutants (I50N and G51T) with null AP functional scores (FS=0.11 and 0.08 respectively) for their ability to auto-cleave in the bacterial assay as above. We found that I50N and G51T PR were as defective in AP as the active site mutant D25A and were completely unable to cut themselves out of the pre-protease construct (Fig. 2F).

In prior work, we measured the effects of PR mutations on viral fitness by individually randomizing each amino acid position of PR in the NL4-3 strain of HIV-1 and quantifying the experimental effects of each mutation during infection of a T-cell line (45). Because PR not only needs to cut itself out of the Gag-pol polyprotein but subsequently must cleave 10 additional cut-sites in the process of generating infectious viruses, viral fitness measurements integrate the cutting function of both mature and precursor protease. A defect in any critical protease reaction will cause a defect in fitness. Based on the viral fitness scan, the WT amino acid is critical for fitness at 51 positions of PR, defined as 15 or more mutations having null-like function at these positions (Fig 2G). These 51 fitness critical positions are located throughout the PR structure, in contrast to the 23 critical positions for AP function which cluster at the active site, dimerization site, and flaps. We reason that positions that are critical for fitness, but not AP are primarily required for cutting by mature protease. Broader mutational constraints for mature protease compared to AP are consistent with the defined structure of mature protease, which places more positions in defined physical environments that in turn restricts the mutations that are compatible with structure and function.

To examine similarities and distinctions between the mechanism of AP and cutting by mature protease, we compared mutational scanning results with our yeast AP assay and our previous analysis of viral fitness (Fig. 3). To facilitate identification of separation of function mutations (mutations that support AP but are deleterious for fitness) we used cutoff scores geared to include most variants that were functional for AP and strongly deleterious for viral fitness (pink shaded quadrant in Fig. 3A). Like all classification approaches, this approach is not perfect. Of note, our cutoffs were chosen to minimize false positive separation of function variants. Overall, about 85% of variants that we classify as non-functional for AP are strongly deleterious for viral fitness (Fig. S1B, peak 3), indicating that the cutoffs generally capture the requirement of AP for fitness.

**Figure 3.**
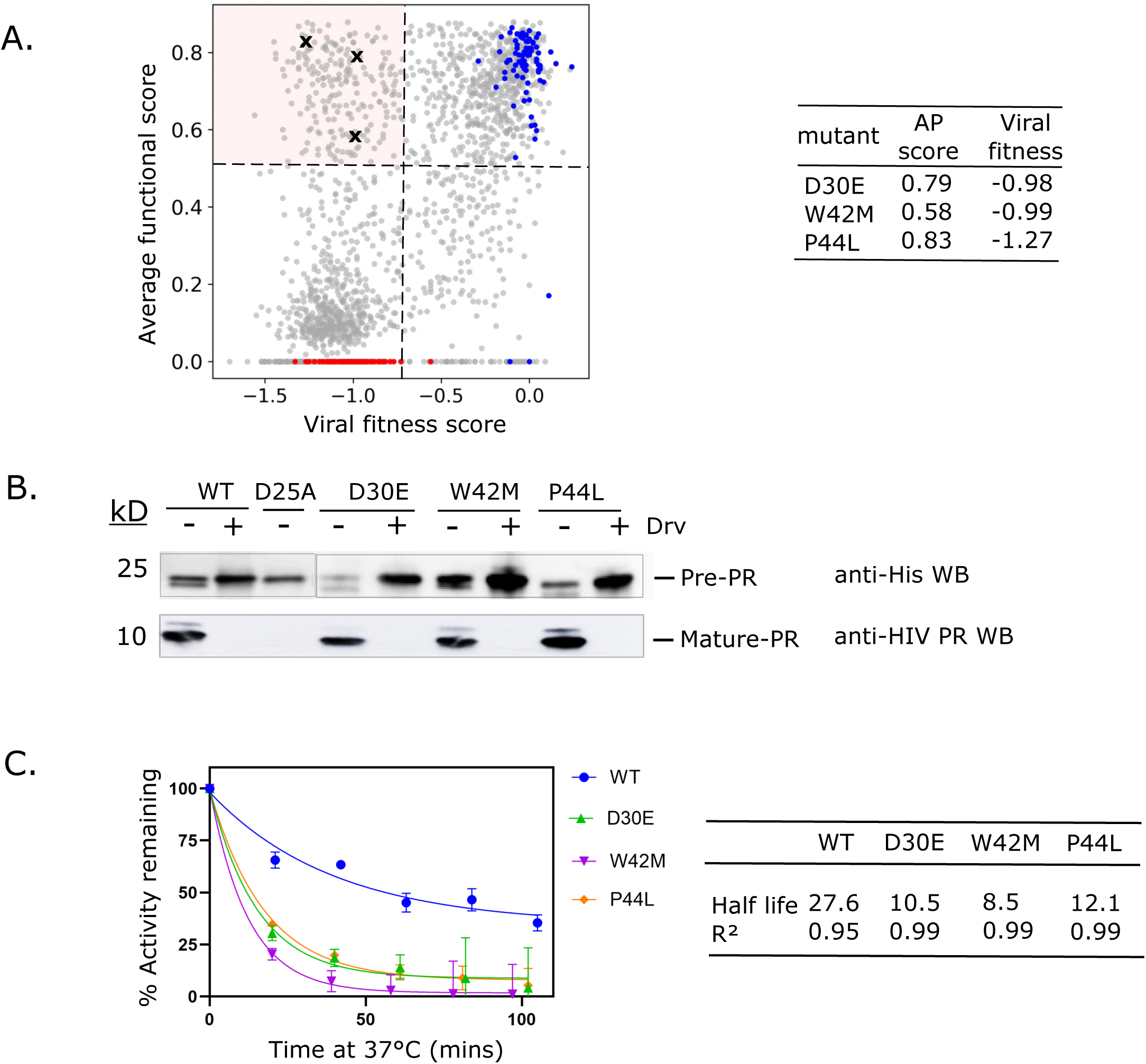
Identification of separation of functions mutations that support AP but are deleterious for fitness. A) Comparison of functional scores from AP assay to viral fitness score (45) for each mutant in PR library. Dashed lines indicate 4 standard of deviations below the distribution of WT synonyms. Stop codons are shown in red and wild-type synonyms in blue. Pink quadrant signifies separation of function mutants. B) Three separation of function mutants (marked with X in panel A) were measured independently for AP activity by their ability to cut themselves out of minimal constructs when expressed in bacteria. All three variants efficiently cut themselves out of minimal constructs as visualized by the disappearance of pre-protease and appearance of mature protease in a Drv-dependent fashion. Three replicates were analyzed and showed similar results. C) The three identified separation of function mutants were cloned and purified as mature PR variants. The PR variants were incubated at 37 °C for the indicated amount of time at which point their enzymatic activity was measured using a peptide substrate at 37°C. Half-lives of PR variants at 37 °C were estimated by fitting the decay curves to an exponential equation.

Among the variants we identified as functional for AP, 222 (25%) were strongly deleterious for fitness and we considered these separation of function mutations as likely defective for cleavage as mature protease. We independently investigated three of the separation of function mutations in isolation (D30E, W42M and P44L) for AP and cleavage of peptide substrates in mature form (marked with Xs in Fig. 3A). We assessed AP by measuring the ability of the variants to efficiently cut themselves out of minimal constructs when expressed in bacteria. All three separation of function variants efficiently cut themselves out of minimal constructs as visualized by disappearance of pre-protease and appearance of mature protease in a Drv-dependent fashion (Fig. 3B), consistent with their AP function in our yeast screen. Additionally, as purified mature protease proteins, all three separation of function variants exhibited strong defects in peptide cleavage assays at physiological temperature (Fig. 3C), providing a rationale for their defects in viral fitness. These detailed studies support our interpretation of the high-throughput studies and support our identification of a large group of mutations that maintain auto-cleavage function but are unable to efficiently cleave substrates as a mature protease.

Having confirmed the separation of function properties of these individual variants by independent assays, we sought to understand how they might influence protease structure and dynamics. Separation of function mutations occurred throughout primary structure (Fig. 4A) but tended to cluster at hot-spot positions (Fig. 4B&C). Clustering at hot-spot positions is consistent with the idea that positions that increase in dynamics during AP compared to mature PR would show decreased mutational sensitivity.

**Figure 4.**
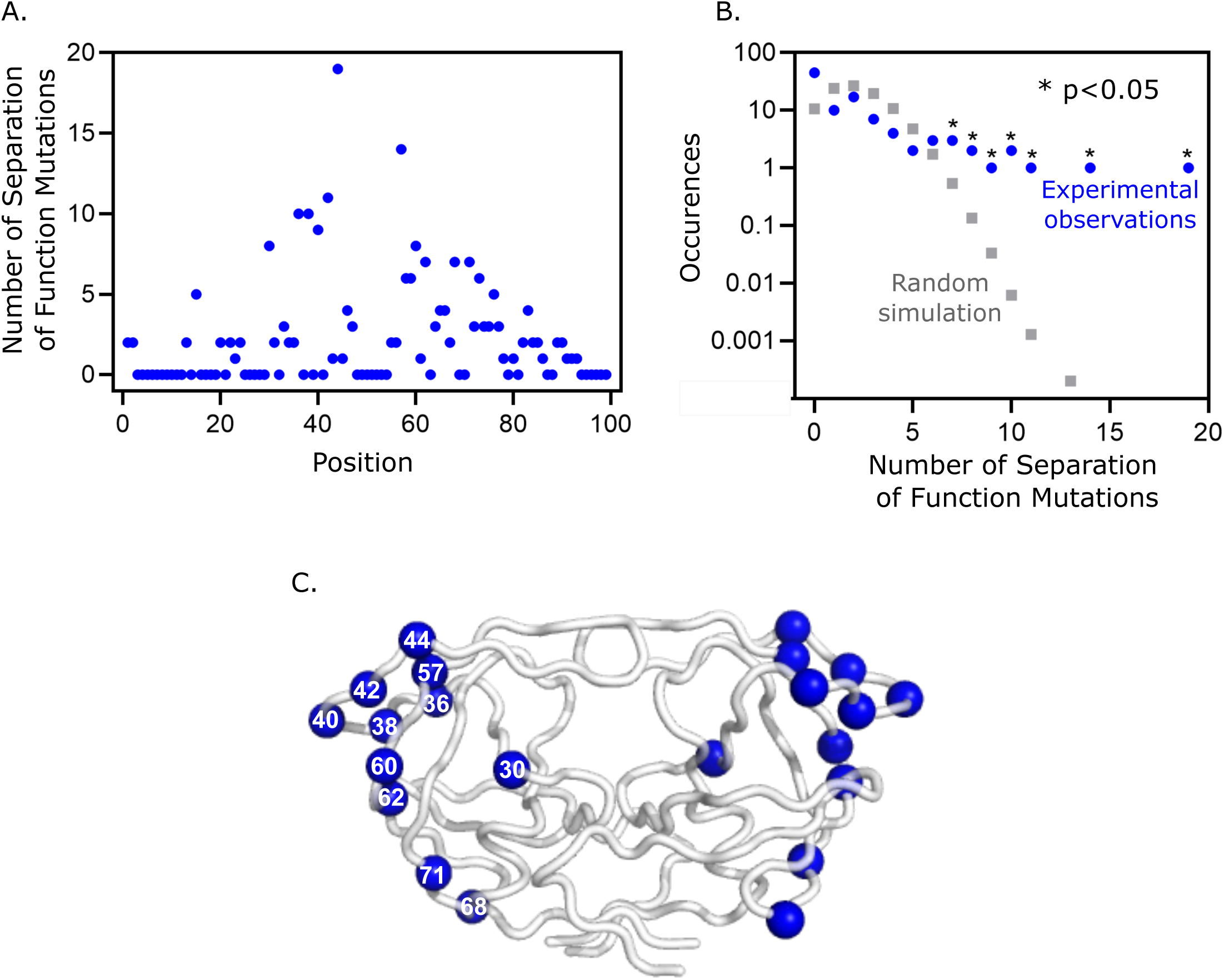
Structural distribution of PR separation of function mutations. A) The number of identified separation of function mutations at each position of PR. B) Clustering of variants identified as separation of function mutants at positions in PR compared to random sampling. C) Separation of function mutations tended to cluster at hot-spot positions. Positions with 8 or more separation of function mutations are shown on both PR subunits as blue spheres.

Because the structure of protease during AP shows more dynamics than mature PR, we reasoned that regions that were unstructured for AP but not mature cutting would harbor separation of function mutations. These separation of function positions would tolerate mutations for AP where they are dynamic, but not for cutting by mature PR (and therefore viral fitness) where they are required for folding of the entire protein. Building on this idea, we performed molecular simulations to provide views of AP mechanism informed by our mutational scan results (Fig. 5). Starting with a structure of mature protease bound to a peptide substrate from the MA-CA cut site (1KJ4.pdb (21)), we modeled the AP cut site connected to the N-terminus of PR into the active site. We then used the program Modeller(46, 47) to simulate physically realistic unfolded conformations for regions that were mutationally tolerant for AP function (positions that tolerated 17 or more mutations for AP) or that were hot-spot positions for separation of function (Fig. 4). We generated and overlayed 10 of these models to provide a rough sense of the dynamic structure of protease during AP compared to the mature structure (Fig 5A&B). Importantly, these simulations are meant to indicate the regions of protease that are likely dynamic based on the mutational data and not the fine details of the motions of protease that would require in depth molecular dynamic simulations including other sections of the polyprotein and are beyond the scope of this work. Of note, the regions of protease that are critical for AP based on mutational analysis demarcate a small structure that our data suggests is the minimal stable conformation required for AP function (Figure 5C).

**Figure 5.**
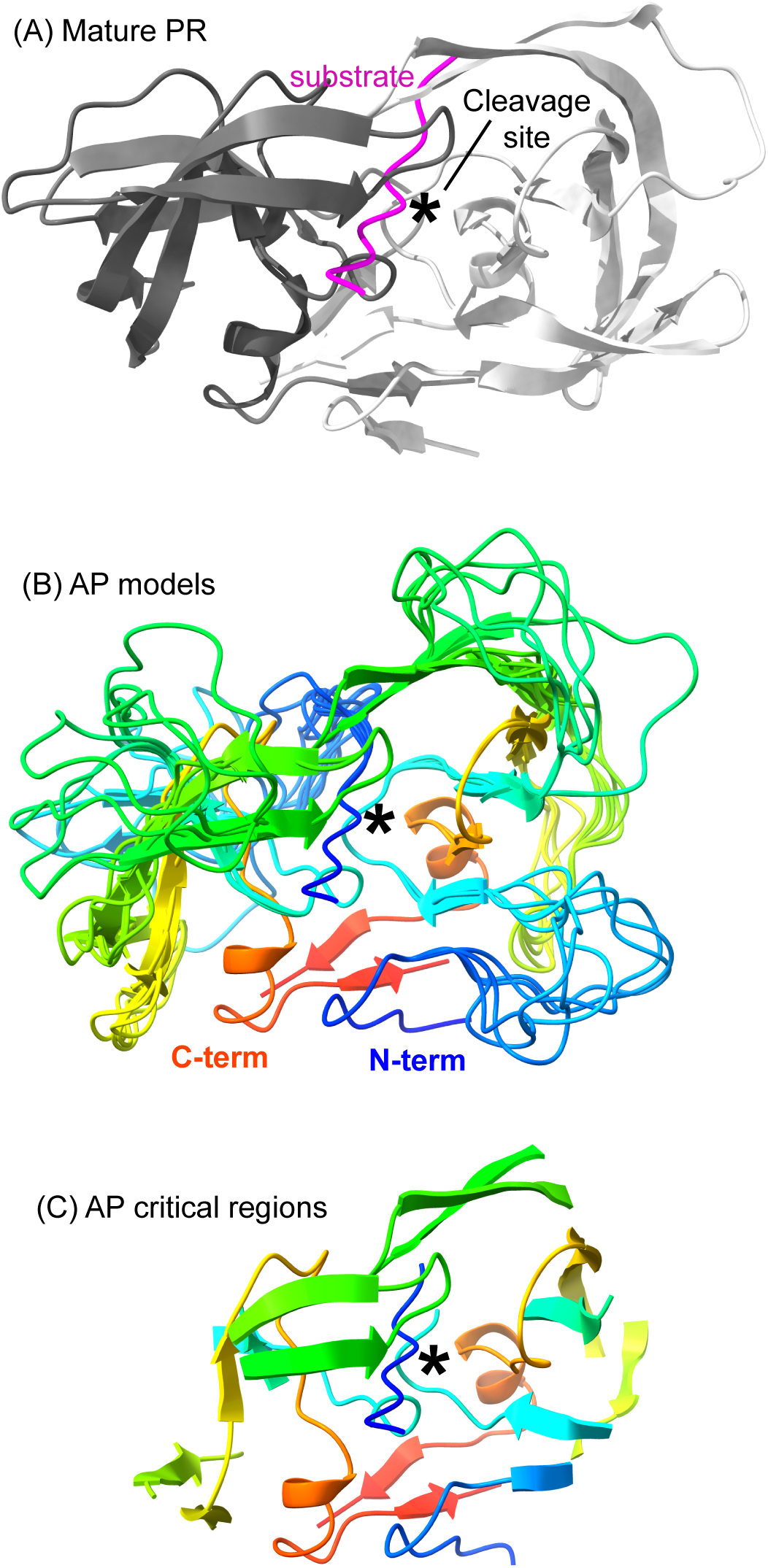
Mutation-informed structural models of AP. A) Molecular representation of mature protease bound to peptide substrate based on 1KJ4.PDB (21). B) Representation of 10 models where regions that were classified as dispensable for AP were modeled as unstructured loops. Structure is colored by rainbow spectrum beginning with blue for the N-terminus and ending with red for the C-terminus C) Molecular representation of regions containing positions critical for AP. Our mutational data suggests that this is the minimal structure required for AP.

## Conclusions

Separation of function mutations that support AP, but not viral expansion indicate that there are fewer sequence constraints for AP compared to fitness, which also requires cutting of other sites in the polyprotein by mature protease. These observations are consistent with the high degree of structural dynamics for PR in the polyprotein compared to in its mature form (34) and the underlying principle that ordered structures impose greater mutational constraints than disordered conformations(48). In stable structures, most positions are subject to specific chemical environments that strongly favor the physical properties of specific amino acids. This logic provides a framework for interpreting our mutational data on AP in terms of structure and mechanism.

Our mutational scan of AP provides novel information on sequence constraints that provides a new view of structural dynamics and mechanism. Molecular simulations informed by our mutational data provide a novel view of the highly dynamic process of AP carried out by HIV-1 PR. Our structural models of AP suggest new strategies for designing inhibitors with improved efficacy towards AP. The ensemble of structural models (Fig. 5) provides a data-informed view of the regions of PR that are consistently structured and those that are highly dynamic during AP. Inhibitors that target regions that are consistently structured during AP should result in improved efficacy compared to regions that are highly mobile (because favorable enthalpic contacts in these regions would be counteracted by unfavorable entropic contributions from reduced motions). The new views of AP provided here provide important guides for understanding and potentially inhibiting a critical step in the infection cycle of HIV. AP is critical to a wide range of viruses of pandemic concern including SARS-CoV-2, HCV, Dengue, West Nile Virus, etc (49). Our approach is general and can be directly applied to understand the critical, but enigmatic process of AP in many pathogenic viruses.

## Materials and Methods

### Generating the yeast reporter strain and plasmids

We generated the DBY646 yeast reporter strain (W303 *HO::Gal1p-GFP,His3; Gal4::Trp1; Gal80::Leu2; Pdr5::NatMX4*) by making genomic changes to the W303 strain using standard yeast genetic manipulation techniques (50, 51). The GFP reporter was integrated at the HO locus and engineered with 500 bases of the Gal1 promoter that can be activated by the Gal4 transcription factor. Disruption of Gal4 and Gal80 were performed to eliminate the ability of the yeast to alter their metabolism in the presence of engineered Gal4 transcription factors. All genomic changes were targeted using at least 45 bases of homology at either end of linear DNA constructs and confirmed by PCR using primers that span across both newly formed junctions in the engineered strain.

The split Gal4 transcription factor with PR in the middle (Gal4 DBD-PR-AD) was generated by overlapping PCR as previously described (44). The resulting hybrid protein is composed of the DNA binding domain of Gal4 (aa 1-147), the HIV-1 pol gene fragment encoding 61 aa upstream of PR, PR (99 aa), and 25 aa downstream of PR, and the activation domain of Gal4 (aa 768-881) (termed Gal4 DBD-PR-AD). The protease sequence is derived from the HIV-1 NL4-3 strain. The resulting Gal4 DBD-PR-AD fusion gene was cloned by Gibson assembly downstream of a tetracycline responsive element on a p417 plasmid containing a KanMX marker. Additionally, a gene encoding the tTA protein was cloned into this vector downstream of a CMV promoter. The tTA gene enables tight repression of the tetracycline-driven promoters in the presence of drug. The mCherry gene was subsequently cloned downstream of a TEF promoter to create the plasmid 417 tetO2-Gal4-DBD-PR-AD (WT); CMV-tTA; TEF-mCherry (pDB664). A destination vector for the library was generated by removing the PR sequence and replacing it with a restriction site for KpnI. The D25A and I13L point mutations were generated by site-directed mutagenesis.

### Generating comprehensive point mutant library in the self-cleavage reporter

Point mutation libraries were generated using a cassette ligation strategy as described (Hietpas et al). In short, the Gal4 DBD-PR-AD fusion gene was inserted into a 417 shuttle plasmid by Gibson assembly. Primers were designed for whole-plasmid PCR to generate vectors with inverted type IIS BsaI restriction sites. Annealed oligonucleotide cassettes with a single codon randomized as NNN and ends complementary to the 5’ BsaI overhangs of the vector were ligated into the vector and transformed into *E.coli*. Transformants were grown in mixed liquid culture from which plasmid DNA was isolated.

The resulting library was combined via Gibson assembly with the destination vector described above. To avoid bottlenecking the library, sufficient transformations were performed to recover more than 50 independent transformants for each PR variant in the library. To improve accuracy of deep sequencing steps, each variant of the library was tagged with a unique barcode. A pool of DNA constructs containing a randomized 18 bp barcode sequence (N18) was cloned into BsrDI and EcoRI sites upstream of the tet promoter via restriction digestion, ligation and transformation into chemically competent *E.coli*. These experiments were designed to have each PR variant represented by 10-20 unique barcodes. To associate barcodes with PR variants, we performed paired-end sequencing using a primer that reads the N18 barcode in Read 1 and a primer that anneals downstream of the protease in Read 2. To facilitate efficient Illumina sequencing, we created shorter PCR products by removing the region between the barcode and the PR library by restriction digest with EcoRI and BglII, followed by blunt ending with T4 DNA polymerase and plasmid ligation at a low concentration. The resulting DNA was relinearized by restriction digest with AscI and XhoI and amplified with 8 cycles of PCR to generate products for Illumina sequencing. The resulting PCR products were sequenced using a MiSeq instrument with asymmetric reads of 50 bases for Read 1 (barcode) and 250 bases for Read 2 (PR sequence). After filtering low-quality reads (Phred scores < 10), the data was organized by barcode sequence. For each barcode that was read more than three times, we generated a consensus of the PR sequence that we compared to wild type to call mutations.

### Yeast competitions and flow cytometry

The plasmid library was transformed using the lithium acetate procedure into the DBY646 reporter strain described above, ensuring greater than 10-fold transformation coverage of each barcoded PR variant. Following 12 hr of recovery in SD-H-L-W + 2 µg/mL doxycycline at 30°C, transformed cells were washed three times with SD-H-L-W + 2 µg/mL doxycycline + 0.4 mg/mL G418 (SD-H-L-W+doxy+G418) to remove the extracellular DNA and grown in SD-H-L-W+doxy+G418 at 180 rpm at 30°C for 24 hours. This library was supplemented with 20% glycerol, aliquoted and frozen at −80°C.

An aliquot of frozen yeast cells transformed with the library was thawed and amplified in 75 mL SD-H-L-W+doxy+G418 overnight at 30°C. Subsequently, the library was diluted to early log phase in 100 mL of SD-H-L-W+doxy+G418 and grown for 8 hours after which the culture was split in half, spun down and resuspended in SD-H-L-W+G418 with either 0 or 30 µM Darunavir. Doxycycline was omitted to induce expression of the PR construct. Cultures were grown with shaking at 180 rpm for 12 hours at which point samples were collected for FACS analysis.

### FACS sorting

Cells were washed two times with tris-buffered saline containing 0.1% Tween and 0.1% bovine serum albumin (TBST-BSA). Cells were diluted to 10^6^/mL and sorted based on GFP and mCherry expression on a FACS Aria cell sorter into cells expressing cut transcription factor (low GFP expression) in one population and uncut transcription factor (high GFP expression) in a second population. Collected cells were outgrown in 50 mL SD-H-L-W+doxy+G418 media overnight at 30°C following which cell pellets were spun down and stored at −80°C.

### DNA preparation and sequencing

We isolated plasmid DNA from each FACS cell population as described(52). Purified plasmid DNA was linearized by AscI. Barcodes were amplified with 20 cycles of PCR using Phusion polymerase and primers that add Illumina adapter sequences and a 6 bp identifier sequence used to distinguish cell populations. PCR products were purified twice over silica columns and quantified using the KAPA SYBR FAST qPCR Master Mix (Kapa Biosystems) on a Bio-Rad CFX machine. Samples were sequenced on an Illumina NextSeq instrument in single end 75 bp mode.

### Analysis of Illumina sequencing data

We analyzed the Illumina barcode reads using custom scripts that have been deposited on GitHub (https://github.com). Illumina sequence reads were filtered for Phred scores>10 and strict matching of the sequence to the expected template and identifier sequence. Reads that passed these filters were parsed based on the identifier sequence. For each screen/cell population, each unique N18 read was counted. The unique N18 count file was then used to identify the frequency of each mutant using the variant-barcode association table. To generate a cumulative count for each codon and amino acid variant in the library, the counts of each associated barcode were summed.

### Calculation of functional effects from sequencing data

To determine the functional score for each variant in the FACS screen, the fraction of each variant in the cut and uncut windows was first calculated by dividing the sequencing counts of each variant in a window by the total counts in that window. The functional score was then calculated as the fraction of the variant in the cut window divided by the sum of the fraction of the variant in the cut and uncut windows. Variants that result in a fully cut TF, in addition to variants that encode for a stop resulting in a truncated TF, will have low GFP signal and fall in the cut window, resulting in a functional score of 1.

### Generation of plasmid expressing the PR precursor

The PR precursor sequence containing 59 amino acids upstream and 23 amino acids downstream of the protease was transferred from the pDB664 plasmid described above to a pET21 vector, downstream of the T7 promotor. The Q7K mutation was generated by site-directed mutagenesis in the PR sequence to increase stability and reduce intramolecular autoproteolysis of the protease (53). Additional individual point mutants were generated in this construct by site-directed mutagenesis and confirmed by Sanger sequencing

### Partial purification and analysis of PR precursor construct by SDS-PAGE

Autoprocessing of the PR precursor was monitored as previously described (35) with the following modifications. Cultures of *E.coli* BL-21 (DE3) pRIL harboring the pET21 PR precursor plasmids were grown overnight in Luria–Bertani medium (LB) containing 100 μg∕mL of carbenicillin either with or without 30 µM Darunavir (DrV). Cells were diluted to an optical density at 600 nm (OD_600_) of 0.15 in the same media with or without DrV and then grown for 2 hours at 37 °C to an OD_600_ of ∼0.8 at which point protein expression was induced by the addition of 2 mM isopropylthiogalactoside (IPTG) for one hour. Cells were harvested and pellets were stored at −20°C. The expressed protein was partially purified as follows: Frozen pellets were thawed, resuspended in 0.5 mL 50 mM Tris-HCl, pH 8, 10 mM EDTA and 5 mM DTT (buffer A), and lysed by sonication in the presence of 100 μg∕ mL lysozyme. The insoluble fraction was recovered by centrifugation in an Eppendorf centrifuge spun at 15,000 rpm for 20 min at 4 °C. The pellet was suspended by brief sonication in 0.5 ml of 50 mM Tris-HCl, pH 8, 10 mM EDTA and 5 mM DTT containing 1 M urea and 0.5% Triton X-100 and recovered again by centrifugation. The final pellet was rinsed in buffer A and then dissolved in 50 µl SDS-PAGE sample buffer and boiled for 5 minutes. Samples were centrifuged and analyzed by SDS-PAGE (4-20% gradient Bio-Rad gel). Proteins were visualized by staining with SYPRO Orange (Invitrogen). The density of gel bands was quantified by an Amersham Imager 600 and analyzed by the software provided by the manufacturer.

### Analysis of autoprocessing of precursor PR by Western blot

The partially purified lysate from above was diluted 1:100 and analyzed via SDS-PAGE electrophoresis and western blotting. The PVDF membrane was blocked in 5% milk and TBS at room temperature for 1 hour. After blocking, the membrane was either probed with primary antibody for anti-HIV PR (MA1-19015; recognizes free N-term of PR), or anti-His primary antibody (AD1.1.10; BioRad) at a 1:3000 dilution in 5% milk and TBS overnight at 4°C. After washing in 1xTBS, the membrane was incubated in anti-mouse secondary conjugated to HRP at a 1:5000 dilution in 4% milk and TBS for 1 hour at room temperature. The surface of the membrane was incubated in ECL substrate for 5 minutes. Bands were visualized using chemiluminescence. β-galactosidase reporter activity of yeast lysates was quantified as described (54).

### Generation of HIV-1 PR variants, protein expression and purification

Mature HIV-1 PR variants were isolated by PCR from the pET21 precursor PR plasmid and cloned into the bacterial expression vector, pET11 (Novagen). PR sequences were confirmed by Sanger sequencing. The bacterial expression and purification of all HIV-1 PR variants were carried out essentially as previously described(55). Briefly, the plasmid encoding HIV-1 PR was transformed into BL-21(DE3) pRIL bacterial cells. Cells were grown at 37^ᵒ^C in LB to an OD_600_ of 1.0, protein expression was induced by the addition of 2 mM IPTG for one hour, and harvested by centrifugation. Following lysis with a cell disruptor, HIV-1 PR was purified from inclusion bodies. The inclusion body pellet was dissolved in 50% acetic acid and HIV-1 PR was further purified using a 100 ml Sephadex G-75 superfine column (Sigma Chemical) equilibrated with 50% acetic acid. Fractions containing HIV-1 PR were combined and refolded by 10-fold dilution into ice cold refolding buffer (0.05 M sodium acetate, pH 5.5, 5% ethylene glycol, 10% glycerol, 5 mM DTT). Purified HIV-1 PR was concentrated using Amicon-10 concentrators and stored at −80^ᵒ^C. The total protein concentration for each variant protease dimer was measured by absorption at 280 nm using an extinction coefficient of 24,980 M^-1^cm^-1^.

### FRET assay to measure enzymatic activity of HIV-1 PR variants

To determine the enzymatic activity of each HIV-1 PR variant a Förster Resonance Energy Transfer (FRET) assay was performed as previously described (56). In this assay, the PR substrates consist of an eight amino acid cleavage site of MA/CA in gag polypeptide: SQNY/PIVQ with a fluorescent donor, 5-[(2-aminoethyl) amino] naphalene-1-sulfonic acid (EDANS), attached at the N-terminus and a quenching acceptor, 4-(4-dimethylaminophenylazo)benzoic acid (DABCYL), attached at the C-terminus. The fluorescence of EDANS is low in these substrates due to quenching by DABCYL. EDANS fluorescence is increased following cleavage of the peptides and separation from DABCYL, and thus the rate of cleavage can be measured by the increase in fluorescence emission at 492 nm over time. The assay was performed at 37°C. MA/CA peptide was dissolved to a final concentration of 20 µM in 4% DMSO. 5 nM HIV-1 PR in 50 mM sodium acetate, pH 5.5, 100 mM NaCl was added after incubation at 37°C for the indicated time. The change in fluorescence over time was measured in a fluorimeter with an excitation at 340 nm and emission at 492 nm and monitored for 15 minutes. The initial rate fluorescence increase was measured. Protease enzymes were incubated at 37°C and activity was measured at indicated time points by dilution of enzyme and substrate into prewarmed buffer in a prewarmed cuvette. Three independent experiments were performed and averaged.

### Molecular modeling

We identified positions that our mutational data indicated were unimportant for AP. We considered both positions that were hot-spots with seven or greater separation of function mutations (required for fitness, but not AP) and positions that tolerated 17 or more different amino acids for AP (functional score > 0.5). In defining dynamic regions we included positions adjacent to those unimportant for AP to enable structural flexibility even for regions with a single position (as long as the adjacent positions were not critical for AP -defined as 17 or more amino acids causing deleterious AP (functional score < 0.5).

We started with the 1KJ4.PDB structure (21) of mature protease bound to a peptide substrate from the MA-CA cut site. We chose this starting structure because of similarities in sequence of the MA-CA cut site and the AP cut site at the beginning of PR. There is no structure of PR bound to the AP cut-sites. The MA-CA site shares many similarities especially adjacent to the scissile bond where they both have an aromatic residue (Tyr or Phe) at the P1 position and Pro at the P1’ position. We mutated the P4 through P4’ positions in the substrate to the AP cut-site identity in Pymol (Schrodinger Inc) choosing side chain geometries that minimized atomic overlaps. We deleted the first four amino acids from the A chain of PR and then connected position 5 in PR to position 4 in the substrate. We used Modeller (47) to generate “loop” conformations of positions 4-11 in the A chain that were physically reasonable. We performed similar Modeller “loop” simulations to generate conformations for regions that we classified as dispensable for AP function. We eliminated “loop” conformations that exhibited strong atomic overlaps with other parts of the structure.

## Acknowledgements

This work was supported in part by grant R21AI157871 from the National Institutes of Health to DNAB.

## Figure Legends

**Figure S1. Some mutations that fully unfold PR lead to non-AP dependent inactive reporter activity.** A) Distribution of Drv responsive score for all variants (gray), stop codons (red), and wild-type synonyms (blue). The Drv responsive score for each variant was calculated by the functional score in the absence of Drv divided by the functional score in the presence of Drv. B) The distribution of functional effects of all PR library variants (see Fig 2C) has three main peaks. The variants in each peak were analyzed for their ability to support viral fitness based on their viral fitness score(45) (cutoff = −0.75). C) The variants in each peak of the DFE in Fig. 2C were analyzed for their responsiveness to Drv. D) Two mutants with functional scores in the range of stops were measured independently for their ability to cut themselves out of minimal constructs when expressed in bacteria and compared to the catalytically dead variant, D25A.

